# Can We Really See the World in Three Dimensions? An Exploratory Study with Finger Alignment Task and Ponzo Illusion

**DOI:** 10.64898/2026.10.18.759971

**Authors:** Amrita Mukherjee, Avijit Paul, Kuntal Ghosh

## Abstract

Stereopsis is the perception of depth when two 2-dimensional (2D) images of an object from slightly different angles align with corresponding retinal positions in both eyes. However, when we consider the blind spot (BS), a new set of challenges arises. In binocular vision, one eye’s higher visual areas compensate for the other’s non-receptive region. However, our brain only receives information from one image patch in the BS region, and another visual patch does not contribute to constructing a comprehensive 3-dimensional (3D) representation of the scene. This suggests that the brain adeptly manages this dimensional transition. We aimed to determine the extent to which we perceive 3D across our entire field of view (FoV). We employed the Ponzo illusion, a stimulus relying on visual depth cues (pictorial depth cues), and an eye-tracking technique to assess participants’ accurate fixation and eye movement patterns. The participants attempted to align their index fingers before a chart paper to create a Horopter-like setup, supporting the hypothesis about our capacity for stereoscopic perception. The results of our exploratory pilot study with two specific tasks revealed that our ability to perceive depth diminishes beyond approximately 8° of retinal eccentricity (in the horizontal meridian) in both halves of the visual field centered around the fovea. This limitation is not apparent in our day-to-day experience.

## 1 Introduction

When perceiving real-life objects, the distribution of their two-dimensional (2D) Grossberg and Todorovic (1988) spatial information falls onto different angular locations on the retinas Masland (2001) of our two eyes. Conventional understanding suggests that if these locations, known as corresponding points Wheatstone (1838), share the same visual angle Navarro et al. (1993) in both eyes, the resulting images generate a three-dimensional (3D) perception Wang et al. (2025). However, a question arises when we consider the physicality of our blind spot (BS) Ramachandran (1992). Each human eye possesses a BS, devoid of any receptors Mukherjee et al. (2024), where the optic nerve Selhorst and Chen (2009) exits to relay retinal information to the visual cortex Grill-Spector and Malach (2004). In binocular vision Blake and Wilson (2011), the non-receptive area of one eye is compensated by the corresponding higher visual area of the other eye, avoiding visual oddities. However, if our brain only receives one patch of the image for the BS region, the ocular patch from the other eye may not contribute to constructing a 3D view of that area. Consequently, a transitional crisis from 2D to 3D perception should be experienced. Nevertheless, our real-life experiences challenge this notion, leading us to question how our brain bridges the dimensional gap between 2D and 3D perception.

This research delved into the extent of our 3D visual capabilities across the field of view Wheelwright et al. (2018). The core of our 3D visual experience lies in our perception of depth, intertwined with variations in spatial sizes. Previous valuable research has centered on unraveling how depth perception integrates with the 2D spatial environment. It includes in-depth investigations into depth perception algorithms Parker and Cumming (2001); Banks et al. (2012); Munro and Gerdelan (2009); Ohzawa (1998); Cumming (1997); Welchman (2016); Blake and Wilson (1991); Marr et al. (1991); Mayhew and Longuet-Higgins (1982); Fahmy et al. (2013); Vishwanath (2014) and explorations of the da Vinci stereopsis Nakayama and Shimojo (1990). Computational methodologies, such as artificial intelligence/machine learning and neural networks Campesato (2020), have also been harnessed to replicate depth perception Henkel (1997); Pan et al. (2017); Touzene and Larabi (2010); Luo et al. (2016); Zhou et al. (2017). Nevertheless, a significant gap in knowledge remains concerning our ability to perceive depth across our entire field of view Wheelwright et al. (2018).

To address this gap, we conducted two experiments aiming to understand the impact of the cortical representation of real or simulated 3D objects at various retinal eccentricities Navarro et al. (1993) on depth perception in binocular vision Blake and Wilson (2011). In the first experiment, we designed an experimental Horopter Turski (2016) using chart paper, while the second experiment employed the Ponzo illusion Brislin (1974) as a stereopsis stimulus and combined it with eye-tracking technology Majaranta and Bulling (2014) to assess gaze fixation Buchan et al. (2007) and saccadic movements Liversedge and Findlay (2000).

Through these experiments, we seek to provide empirical insights into the mechanisms underlying our 3D visual perception and shed light on the intriguing dimension gap between 2D and 3D experiences. By addressing this critical research gap, we aim to advance our understanding of human visual perception and contribute to the development of more comprehensive computational models and neuroscientific theories in 3D vision.

## 2 Materials and Methods

20 subjects (aged between 20 and 40 years with normal or corrected-to-normal visual acuity) naive to the purpose of our experiments participated in the study after providing written consent. We trained the observers to maintain their gaze fixation at particular positions on the chart paper and the computer monitor. They practiced the demo sets 8 to 10 times. The experiments are described in the following sections. We performed 3 trials for both experiments. Experiments were approved by the Ethics Committee-“Review Committee for Protection of Research Risks to Humans” of the Indian Statistical Institute, Kolkata, India. The Ethics Clearance Number is ISI-IEC/2023/01/02.

### 2.1 Experiment 1 with Chart Paper

We drafted equi-angled points on art paper to depict the visual angles and fixed the paper on a wall (Fig.1). The central point is represented as 0°. On both sides, we considered visual angles up to 70° in both the visual halves Wandell et al. (2007). The distance between the subject and the wall was 0.6m. On both sides, between 0° and 8° the points were 2.5° apart and between 10° and 70° they were 5° apart. This experiment was performed both in photopic and mesopic lighting conditions Barbur and Stockman (2010). We measured the luminosity Salpeter (1955) of light using Mavospot2 Ballestrín et al. (2018). The observers used powerless glasses fitted with a laser pointer so that, with the help of the incident laser ray, they could fix their fovea Yuodelis and Hendrickson (1986) properly and precisely. Before starting the experiment, the observers practiced the eye fixation task several times and also practiced keeping their hands steady and straight, for which we attached a lightweight steel chain to the ceiling, as shown in Fig.1. To place the observers’ eyes and the visual angles at the same vertical level, the height of each observer was adjusted according to the height of the chart paper before starting the experimental trial (see the blue dotted line in Fig.1). In the beginning, we instructed the subjects to fixate their eyes at the center (0°) of the paper and to try to align (match) the tips of both their converging index fingers. Afterward, the subjects were instructed to close their eyes and lower their hands to the resting position. A few seconds later, the subjects were again instructed to fixate their eyes at 0° visual angle, and they were asked to perform the aforesaid alignment process at 2.5° in the left visual field, i.e., they were starting to use their peripheral vision to align their fingertips. After that, the subjects were again instructed to close their eyes and move their hands to the resting position. The same process was repeated up to 60° for both the visual halves. The visual angles at which the subjects were able to execute the finger alignment process properly can be thought of as the points at which they achieved appropriate depth perception.

**Figure 1.**
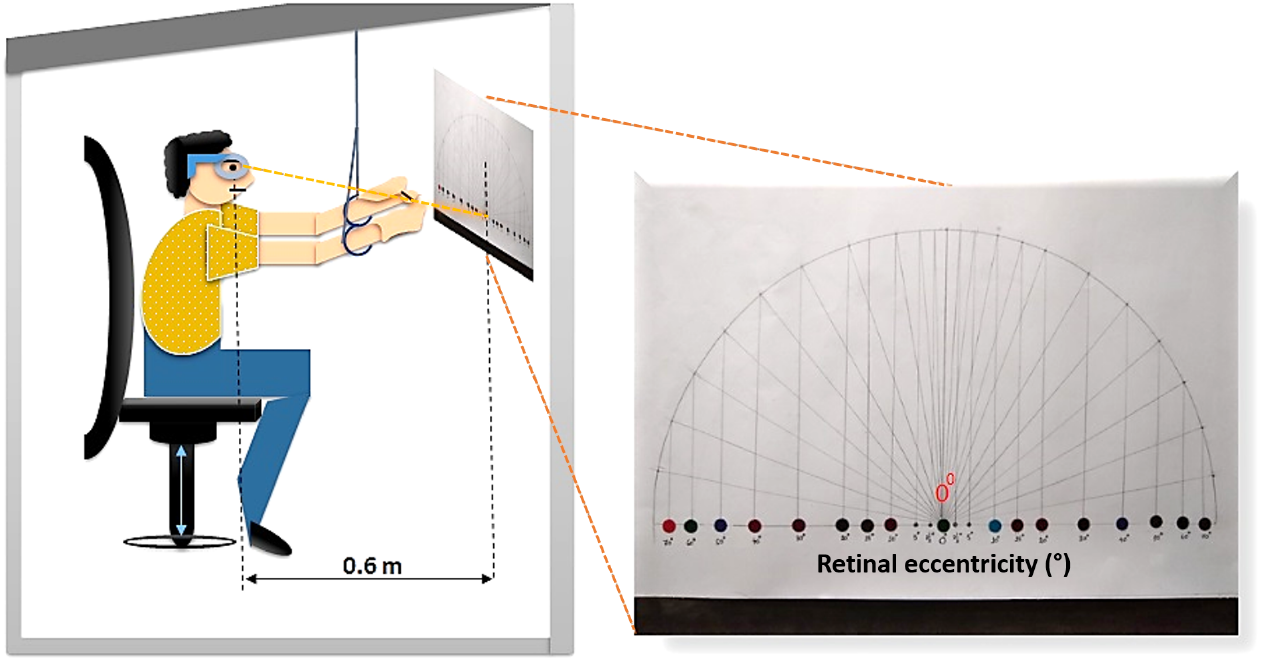
Experimental setup. Snapshot of the Horopter designed on chart paper hanging on the wall. The distance between the observer and the chart paper is 0.6m. The blue dotted lines show the height of the observer’s eye and the chart paper above the ground.

Graphical analysis of task-specific depth perception performance, using data obtained from observers’ fingertip alignment activity under photopic and mesopic lighting conditions, is shown in the graphs (see Figure2). None of the subjects was able to perceive 3D vision beyond a maximum of 8° in either of the left and right visual halves. Some of them could not even align their index fingers beyond 0° retinal eccentricity (see Table 1 for the data analysis).

**Table 1.**
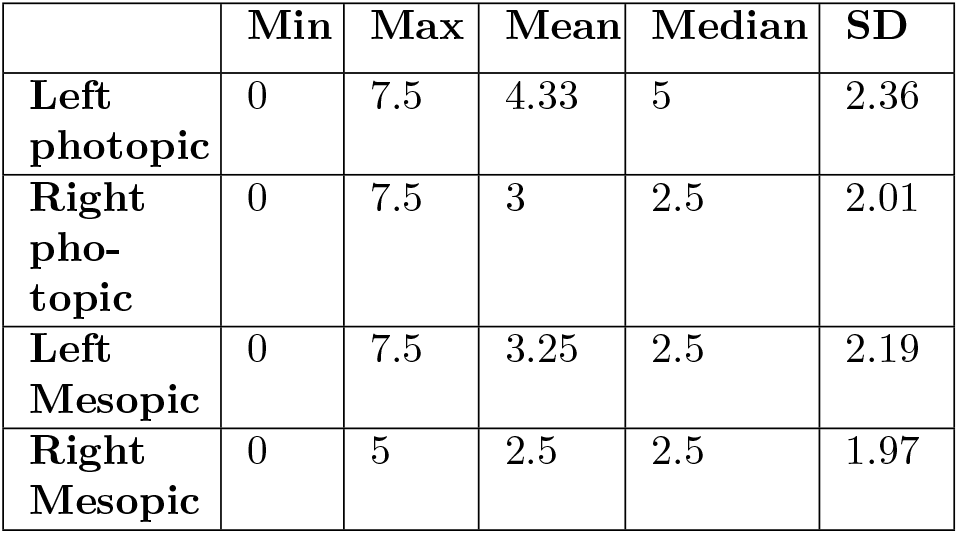
Data Analysis of the task-specific depth perception performance of stereoscopic vision for Experiment 1. The units of all the numerical values are retinal eccentricity (in °)

|  | Min | Max | Mean | Median | SD |
| --- | --- | --- | --- | --- | --- |
| **Left photopic** | 0 | 7.5 | 4.33 | 5 | 2.36 |
| **Right photopic** | 0 | 7.5 | 3 | 2.5 | 2.01 |
| **Left Mesopic** | 0 | 7.5 | 3.25 | 2.5 | 2.19 |
| **Right Mesopic** | 0 | 5 | 2.5 | 2.5 | 1.97 |

**Figure 2.**
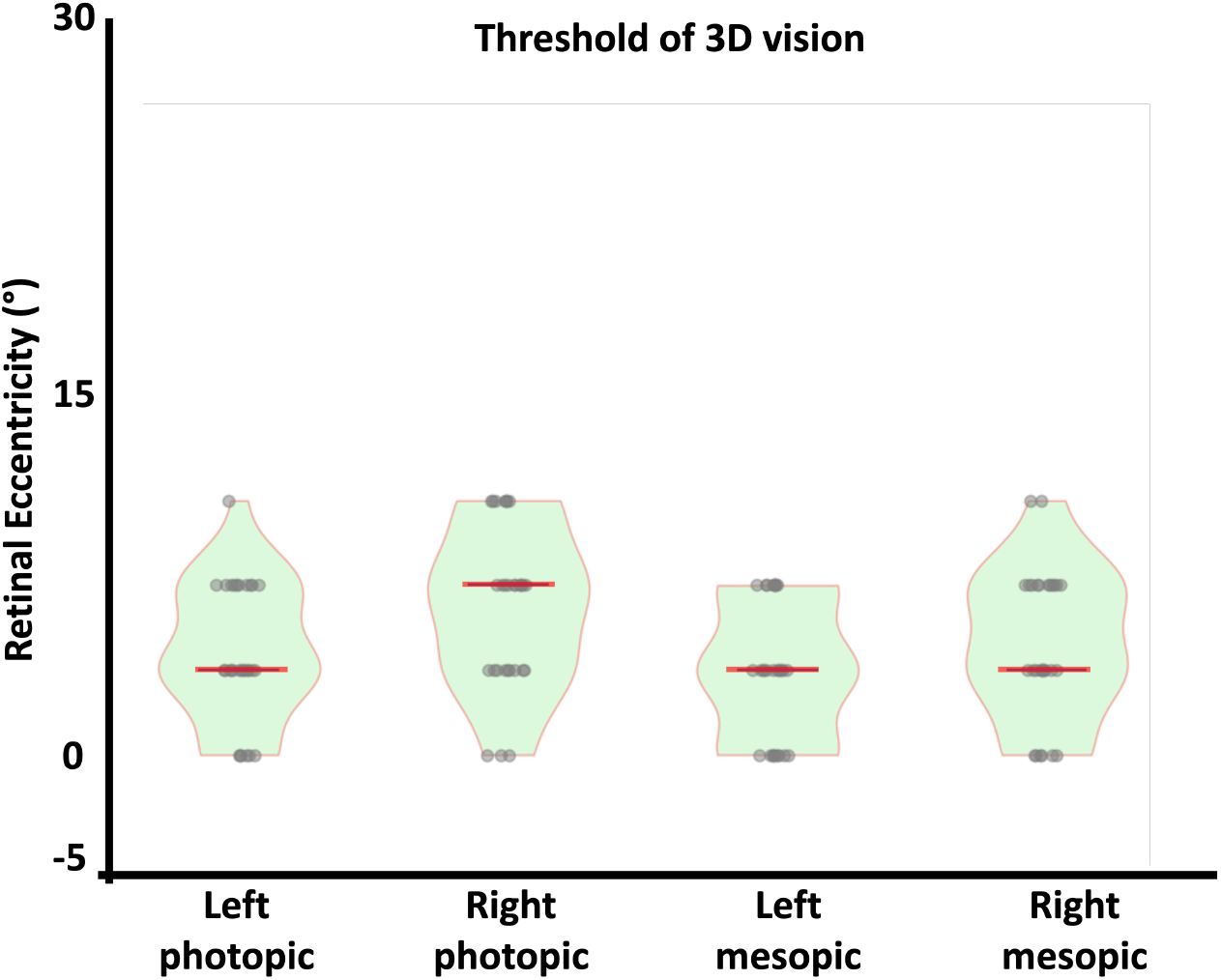
The violin plot of the thresholds of 3D perception (measured in retinal eccentricity) in the left and right visual halves in photopic and mesopic lighting conditions Barbur and Stockman (2010). They are mentioned as Left photopic, Right photopic, Left mesopic, and Right mesopic.

### 2.2 Experiment 2 with Computer Simulation and Eye Tracker

We designed another experiment (this time with computer simulations) to check our depth perception by utilizing a Tobii Pro Nano eye tracker Zhang and Cui (2022) (60Hz sampling frequency, EVT filter Ozun et al. (2010)), and the Ponzo illusion Brislin (1974) as the stimulus. The stimulus was placed at different locations on the computer monitor corresponding to different retinal eccentricities.

First, the artificial stimulus (Fig. 3(a)) was placed at a central position on the screen, and we changed its position along the Left and right directions, starting from the center with a spatial gap of 5 pixels, totaling 40 stimulus positions. We marked a point at the center of the screen for eye-fixation purposes and set the background color to white. We instructed the subjects to fixate on the center point. We asked them to press the mouse anywhere on the monitor if they could perceive the upper bar as longer than the lower one. The position of the stimulus was adjusted in both horizontal directions by pressing any key on the computer. We considered two Areas of Interest (AOI) Hu et al. (2015) - elliptical and rectangular, where the first one was placed around the fixation point, and the latter covered the whole screen. The elliptical AOI was used to check the proper gaze around the fixation point. The rectangular AOI was designed to capture mouse-click events and saccadic movements. We used a single Time of Interest (TOI) Zeng et al. (2026), which is the total recording duration. Thus, only two events were considered: the start and the end point of the TOI. If the subjects observed the upper yellow bar longer than the lower one, they were supposed to experience a depth perception by creating an illusory size difference. We conducted all our experiments under mesopic optical conditions.

**Figure 3.**
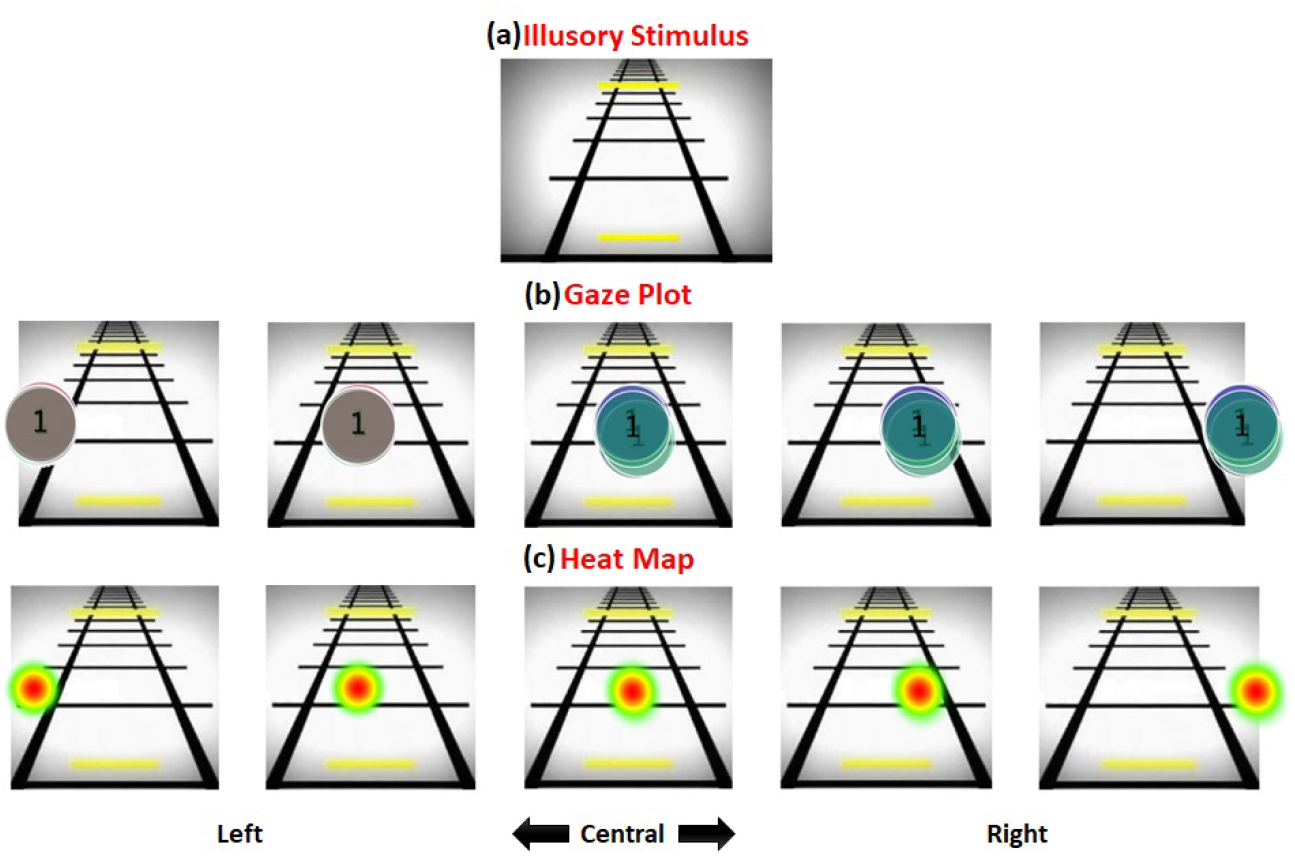
(a) Ponzo illusion Brislin (1974). (b) Gaze plots and (c) heat maps: Though we have taken a total of 40 positions here, we showed gaze plots and the corresponding heat maps for 6 stimulus positions.

We considered various metrics, such as AOI fixation, AOI click, AOI saccade, and Interval, in our experiment. We require the AOI fixation metric for accurate eye fixation. We also use the AOI click metric to identify instances of depth perception. Another important aspect of depth perception is noticing saccadic movements. We designed a rectangular AOI covering the whole screen and considered only the AOI saccade metric, which captures saccadic movements at data acquisition time within the rectangular AOI. Here, we showed the gaze plot Ghose et al. (2020) (Figure 3(b)) and the corresponding heat map Špakov and Miniotas (2007) (Figure 3(c)) for some of the spatial positions. They showed the observers’ appropriate gaze direction and saccadic movements. We analyzed the gaze plots and heat maps because the subjects’ eyes might make a number of rapid saccadic movements to create 3D perception, which could skew the results of our experiment and fail to accurately reflect the subjects’ 3D visual ability. We observed that our subjects performed proper gaze fixation with minimal saccades. This helps us analyze the extent of 3D visual ability for the whole FoV without the effects of saccadic eye movements.

Graphical analysis depicting the retinal eccentricities (we have converted the monitor pixels into visual angles or retinal eccentricities) up to which the subjects reported their opinions about the comparative length of the bars in both left and right horizontal directions is shown in Fig.4. Beyond those limits, they were unable to make out any decision on the comparative lengths. Table 2 describes the data analysis of the stereoscopic vision in the left and right visual halves for both photopic and mesopic lighting conditions Barbur and Stockman (2010) (see Table 2 for the data analysis).

**Table 2.** Data Analysis of the thresholds of stereoscopic vision for Experiment 2. The units of all the numerical values are retinal eccentricity (in °)

|  | Min | Max | Mean | Median | SD |
| --- | --- | --- | --- | --- | --- |
| **Left**<br>**Mesopic** | 2.71 | 4.80 | 3.76 | 3.87 | 0.61 |
| **Right**<br>**Mesopic** | 2.71 | 4.64 | 3.38 | 3.29 | 0.57 |

**Figure 4.**
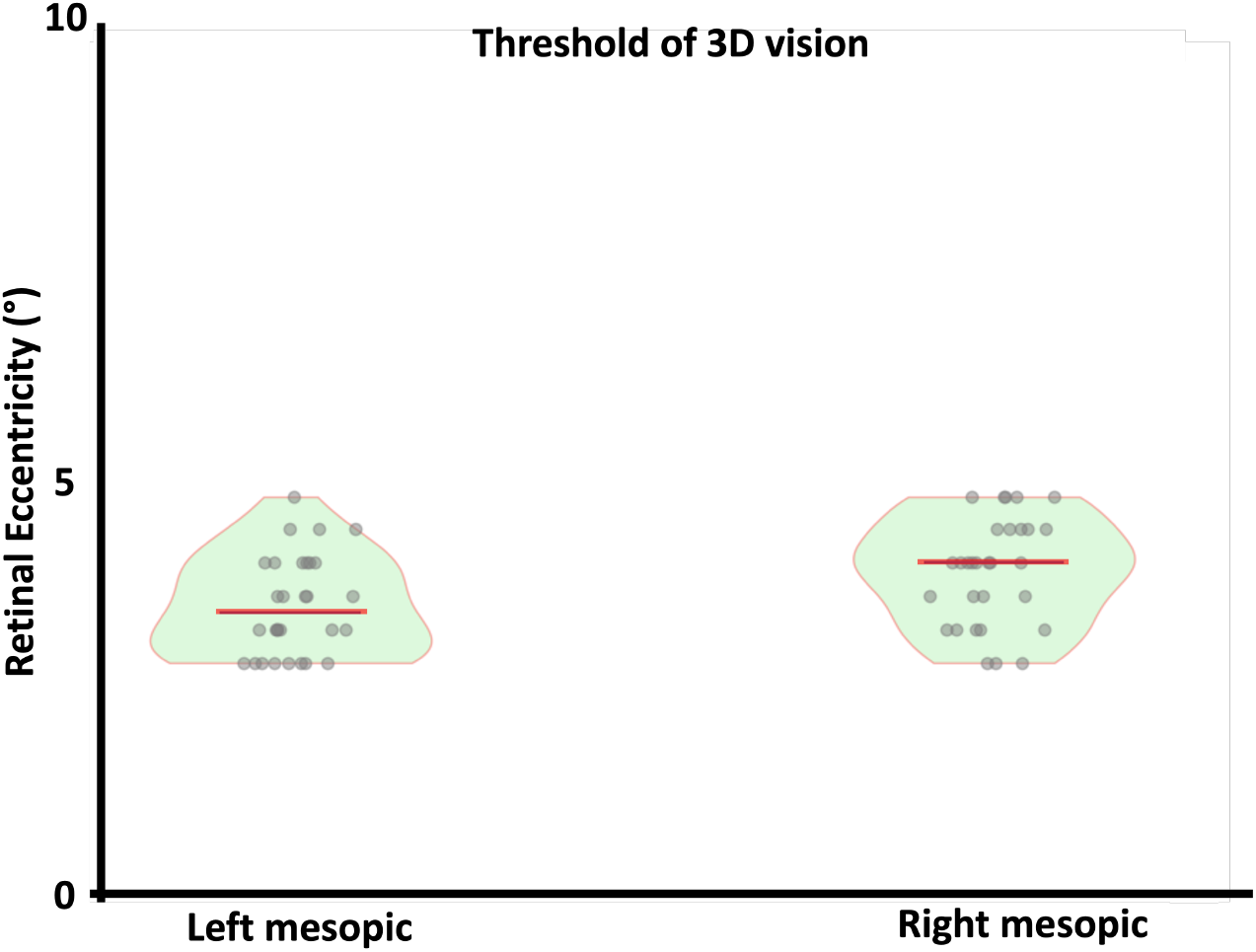
The violin plot of the thresholds of 3D perception (measured in retinal eccentricity) in the left and right visual halves in mesopic lighting conditions. They are labeled as Left mesopic and Right mesopic.

## 3 Discussion and Conclusion

Results of our experiments suggest that the task-specific depth-perception performance of our stereoscopic vision is independent of external lighting conditions or stimulus properties. The findings from both experiments underscore a significant constraint on our ability to perceive 3D visuals, particularly given a maximum retinal eccentricity of 8° in both the left and right visual fields, centered around the fovea. In spite of the fact that the points positioned on the Veith-Müller circle Yeh (1994) generate images on corresponding points of our eyes, our capacity to perceive these points in 3D progressively declines beyond this task-specific depth perception performance, spanning up to approximately 8°. In the initial experiment, participants had difficulty aligning the positions of their index fingertips beyond this angular threshold. Likewise, we noted a similar limitation in the simulated Ponzo illusion experiment Brislin (1974), in which subjects struggled to discern any length disparity between two yellow bars positioned along a virtual rail track.

Conventional belief is that when the visual points falling within the Horopter align with the corresponding points on our eyes, we perceive depth. However, our recent experiment demonstrates a spatial threshold linked to retinal eccentricity, beyond which 3D perception diminishes. Our study indicates that stereoscopic perception is possible within an 8° visual range in both visual halves, starting from the fovea. Notably, retinal eccentricity plays a pivotal role in determining this task-specific depth perception performance, as cone density decreases with increasing eccentricity, resulting in less vivid representation in the visual cortex (i.e., less cortical magnification factor Rovamo and Virsu (1979)). This diminished vividness possibly impacts our ability to perceive depth stereoscopically. We also observed that the left visual half had a slightly higher depth-perception threshold than the right.

Our experiments reconfirm that our perception of the external world is not a straightforward mirror of its objective reality. Instead, it is a cognitive construct shaped by our brains to maximize our survival, albeit within a notably confined 3D framework extending up to approximately 8° in each visual half.

These findings are best interpreted within the present exploratory design and sample, and future work incorporating stereoacuity screening and monocular control would help more precisely isolate the stereoscopic contribution. Because testing focused on the horizontal meridian at a near viewing distance, the results characterize this condition rather than the full field of view. The observed decline may reflect a combination of peripheral acuity, crowding, cortical magnification Rovamo and Virsu (1979), attention, fixation stability, motor precision, and response bias, with cone density as one plausible but untested contributor.

## 4 Acknowledgement

This work is funded by the Cognitive Science Research Initiative (CSRI) of the Department of Science and Technology, Govt. of India, and TAC-DCSW funding of the Indian Statistical Institute. The authors are thankful to all the colleagues in the Laboratory for Cognitive Systems and Cybernetics Research at the Center for Soft Computing Research of the Indian Statistical Institute, Kolkata.

## References

Ballestrín, J., Carra, M. E., Enrique, R., Monterreal, R., Fernández-Reche, J., Polo, J., Casanova, M., Barbero, F. J., and Marzo, A. (2018). Diagnosis of a lambertian target in solar context. Measurement, 119:265–269.

Banks, M. S., Read, J. C., Allison, R. S., and Watt, S. J. (2012). Stereoscopy and the human visual system. SMPTE motion imaging journal, 121(4):24–43.

Barbur, J. and Stockman, A. (2010). Photopic, mesopic and scotopic vision and changes in visual performance. Encyclopedia of the Eye, 3(323-331):2.

Blake, R. and Wilson, H. (2011). Binocular vision. Vision research, 51(7):754–770.

Blake, R. and Wilson, H. R. (1991). Neural models of stereoscopic vision. Trends in Neurosciences, 14(10):445–452.

Brislin, R. W. (1974). The ponzo illusion: Additional cues, age, orientation, and culture. Journal of Cross-cultural Psychology, 5(2):139–161.

Buchan, J. N., Paré, M., and Munhall, K. G. (2007). Spatial statistics of gaze fixations during dynamic face processing. Social Neuroscience, 2(1):1–13.

Campesato, O. (2020). Artificial intelligence, machine learning, and deep learning. De Gruyter.

Cumming, B. (1997). Stereopsis: how the brain sees depth. Current Biology, 7(10):R645–R647.

Fahmy, A. A., Ismail, O., and Al-Janabi, A. (2013). Stereo vision based depth estimation algorithm in uncalibrated rectification. Int J Video Image Process Netw Secur, 13(2):1–8.

Ghose, U., Srinivasan, A. A., Boyce, W. P., Xu, H., and Chng, E. S. (2020). Py-track: An end-to-end analysis toolkit for eye tracking. Behavior research methods, 52(6):2588–2603.

Grill-Spector, K. and Malach, R. (2004). The human visual cortex. Annu. Rev. Neurosci., 27(1):649–677.

Grossberg, S. and Todorovic, D. (1988). Neural dynamics of 1-d and 2-d brightness perception: A unified model of classical and recent phenomena. Perception & psychophysics, 43(3):241–277.

Henkel, R. (1997). A simple and fast neural network approach to stereovision. Advances in Neural Information Processing Systems, 10.

Hu, Y., Gao, S., Janowicz, K., Yu, B., Li, W., and Prasad, S. (2015). Extracting and understanding urban areas of interest using geotagged photos. Computers, Environment and Urban Systems, 54:240–254.

Liversedge, S. P. and Findlay, J. M. (2000). Saccadic eye movements and cognition. Trends in cognitive sciences, 4(1):6–14.

Luo, W., Schwing, A. G., and Urtasun, R. (2016). Efficient deep learning for stereo matching. In Proceedings of the IEEE conference on computer vision and pattern recognition, pages 5695–5703.

Majaranta, P. and Bulling, A. (2014). Eye tracking and eye-based human–computer interaction. In Advances in physiological computing, pages 39–65. Springer.

Marr, D., Poggio, T., Hildreth, E. C., and Grimson, W. E. L. (1991). A computational theory of human stereo vision. In From the retina to the neocortex: selected papers of david marr, pages 263–295. Springer.

Masland, R. H. (2001). The fundamental plan of the retina. Nature neuroscience, 4(9):877–886.

Mayhew, J. and Longuet-Higgins, H. (1982). A computational model of binocular depth perception. Nature, 297(5865):376–378.

Mukherjee, A., Paul, A., Roy, R., and Ghosh, K. (2024). The role of extrinsic and intrinsic factors in perceptual filling-in of the blind-spot with variegated color and texture stimuli. Vision Research, 222:108452.

Munro, P. and Gerdelan, A. P. (2009). Stereo vision computer depth perception. Country United States City University Park Country Code US Post code, 16802.

Nakayama, K. and Shimojo, S. (1990). Da vinci stereopsis: Depth and subjective occluding contours from unpaired image points. Vision research, 30(11):1811–1825.

Navarro, R., Artal, P., and Williams, D. R. (1993). Modulation transfer of the human eye as a function of retinal eccentricity. Journal of the Optical Society of America A, 10(2):201–212.

Ohzawa, I. (1998). Mechanisms of stereoscopic vision: the disparity energy model. Current opinion in neurobiology, 8(4):509–515.

Ozun, A., Cifter, A., and Yılmazer, S. (2010). Filtered extreme-value theory for value-at-risk estimation: evidence from turkey. The Journal of Risk Finance, 11(2):164– 179.

Pan, Z., Jiang, G., Jiang, H., Yu, M., Chen, F., and Zhang, Q. (2017). Stereoscopic image super-resolution method with view incorporation and convolutional neural networks. Applied Sciences, 7(6):526.

Parker, A. J. and Cumming, B. G. (2001). Cortical mechanisms of binocular stereoscopic vision. Progress in brain research, 134:205–216.

Ramachandran, V. S. (1992). Blind spots. Scientific American, 266(5):86–91.

Rovamo, J. and Virsu, V. (1979). An estimation and application of the human cortical magnification factor. Experimental brain research, 37(3):495–510.

Salpeter, E. E. (1955). The luminosity function and stellar evolution. Astrophysical Journal, vol. 121, p. 161, 121:161.

Selhorst, J. B. and Chen, Y. (2009). The optic nerve. In Seminars in neurology, volume 29, pages 029–035. © Thieme Medical Publishers.

Špakov, O. and Miniotas, D. (2007). Visualization of eye gaze data using heat maps. Elektronika ir elektrotechnika, 74(2):55–58.

Touzene, N. B. and Larabi, S. (2010). Disparity map estimation with neural network. In 2010 International Conference on Machine and Web Intelligence, pages 303–306. IEEE.

Turski, J. (2016). On binocular vision: The geometric horopter and cyclopean eye. Vision research, 119:73–81.

Vishwanath, D. (2014). Toward a new theory of stereopsis. Psychological review, 121(2):151.

Wandell, B. A., Dumoulin, S. O., and Brewer, A. A. (2007). Visual field maps in human cortex. Neuron, 56(2):366–383.

Wang, Q., Zhang, Y., Holynski, A., Efros, A. A., and Kanazawa, A. (2025). Continuous 3d perception model with persistent state. In 2025 IEEE/CVF Conference on Computer Vision and Pattern Recognition (CVPR), pages 10510–10522. IEEE.

Welchman, A. E. (2016). The human brain in depth: how we see in 3d. Annual review of vision science, 2:345–376.

Wheatstone, C. (1838). Contributions to the physiology of vision.–part the first. on some remarkable, and hitherto unobserved, phenomena of binocular vision. Philosophical transactions of the Royal Society of London, pages 371–394.

Wheelwright, B., Sulai, Y., Geng, Y., Luanava, S., Choi, S., Gao, W., and Gollier, J. (2018). Field of view: not just a number. Digital Optics for Immersive Displays, 10676:1067604.

Yeh, C.-P. (1994). Depth perception using crossed-looking stereo vision system. In Intelligent Robots and Computer Vision XIII: 3D Vision, Product Inspection, and Active Vision, volume 2354, pages 369–379. SPIE.

Yuodelis, C. and Hendrickson, A. (1986). A qualitative and quantitative analysis of the human fovea during development. Vision research, 26(6):847–855.

Zeng, R., Glaue, E., Moellhoff, N., Giunta, R. E., Lim, T., Felice, F., Prantl, L., Brébant, V., Unbehaun, P., and Frank, K. (2026). How long and how we judge age: A cross-racial and cross-gender eye-tracking investigation. Aesthetic Plastic Surgery, pages 1–9.

Zhang, L. and Cui, H. (2022). Reliability of muse 2 and tobii pro nano at capturing mobile application users’ real-time cognitive workload changes. Frontiers in Neuroscience, 16:1011475.

Zhou, C., Zhang, H., Shen, X., and Jia, J. (2017). Unsupervised learning of stereo matching. In 2017 IEEE International Conference on Computer Vision (ICCV), pages 1576–1584. IEEE.

